# Landscape-scale endophytic community analyses in replicated grapevine stands reveal that dieback disease is not caused by specific fungal communities

**DOI:** 10.1101/2024.04.05.588363

**Authors:** Vinciane Monod, Valérie Hofstetter, Olivier Viret, Vivian Zufferey, Katia Gindro, Daniel Croll

**Affiliations:** Laboratory of Evolutionary Genetics, Institute of Biology, University of Neuchâtel, CH-2000 Neuchâtel, Switzerland; Agroscope, Plant Protection, Mycology, 1260 Nyon, Switzerland; Direction générale de l’agriculture, de la viticulture et des affaires vétérinaires (DGAV), Département de l’Economie, de l’innovation et du sports (DEIS), CH-1110 Morges, Switzerland; Agroscope, Viticulture, Avenue de Rochettaz 21, 1009 Pully, Switzerland

## Abstract

Tree diebacks are complex and multi-factorial diseases with suspected biotic and abiotic components. Microbiome effects on plant health are challenging to assess due to the complexity of fungal and bacterial communities. Grapevine wood dieback is the main threat to sustainable production worldwide and no causality with microbial species has been established. Here, we aimed to test the hypothesis that grapevine esca disease progression has reproducible drivers in the fungal species community. For this, we analyzed a set of 21 vineyards planted simultaneously with a single susceptible cultivar to provide replication at the landscape scale. We sampled a total of 496 plants across vineyards in two different years to perform deep amplicon sequencing analyses of the fungal communities inhabiting grapevine trunks. The communities were highly diverse with a total of 4,129 amplified sequence variants assigned to 697 distinct species. Individual plants varied in fungal community composition depending on the year of sampling, vineyard location, and disease status. However, we detect no specific fungal species driving symptom development across the vineyards contrary to long-standing expectations. Our study shows how landscape-scale replicated field surveys allow for powerful hypothesis-testing for complex dieback disease drivers and prioritize future research towards additional factors.

## Introduction

Dieback is the deterioration of tree health observed increasingly in forests and perennial crops constituting a global concern (Santini *et al*., 2013; Cohen *et al*., 2016; Denman *et al*., 2018). Environmental warming is a key factor in increasing the dieback likelihood and favors the spread of plant diseases (Millar and Stephenson, 2015; Singh *et al*., 2023). Tree diebacks are complex and multi-factorial diseases with biotic and abiotic components (Vieites *et al*., 2009; Tiew *et al*., 2020). Determining the combination of factors leading to decline is challenging (Bettenfeld *et al*., 2020). Vascular wilts are among the most destructive tree declines (Yadeta and J Thomma, 2013). Complex biotic interactions, including polymicrobial and insect activity, influence the onset of dieback and increase severity (Denman *et al*., 2018). Even though most plant diseases are thought to be caused by discrete pathogen species, there is growing evidence that complex plant diseases can arise from synergistic interactions among multiple microorganisms (Lamichhane and Venturi, 2015; Denman *et al*., 2018). Given the complexity of the microbial communities associated with perennial plants, investigating links between microbiome composition and disease status is essential.

Microbiome assembly effects on perennial plant health and dieback rates are challenging to assess due to the large number of coexisting microbes including endophytic fungi (Martins *et al*., 2021). Members of fungal communities interact with each other and with their hosts to cause a wide range of beneficial or pathogenic effects (Compant *et al*., 2019). The microbiota, by providing additional ecological functions to the host (Turner *et al*., 2013), plays a crucial role in plant adaptation to biotic and abiotic environmental conditions potentially enhancing plant health and stress resistance (Pacifico *et al*., 2019; Stewart *et al*., 2021). Microbial communities can promote plant growth by simulating water and nutrient intake, increase health trough antibiosis against pathogens and pests (Rodriguez *et al*., 2009; Rolli *et al*., 2015; Hyde *et al*., 2019; Pacifico *et al*., 2019; Trivedi *et dal.*, 2020). The spectrum of symbiotic associations and their consequences are not well defined and depend on environmental conditions. These associations can transition between commensalism, mutualism or parasitism (Mishra *et al*., 2021). For instance, different strains of *Pantoea ananatis* bacteria isolated from healthy maize seeds have shown diverse effects, ranging from growth promotion to weak pathogenicity or neutral effects on the plant. Such variation in plant-microbe relationships is likely governed by protein secretion systems and effector proteins (Sheibani-Tezerji *et al*., 2015; Stewart *et al*., 2021). Environmental factors and host genotype may also influence the lifestyle of fungi transitioning from endophyte to pathogen as observed in *Fusarium verticillioides* on maize (Bacon *et al*., 2008). Some endophytes display a latent state and turn symptomatic when the plant encounters stress conditions such as drought, humidity, or nutrient starvation (Mishra *et al*., 2021). Fungal endophytes comprise a diverse group of species, some of which are known to cause plant diseases as pathogens, while also being present on asymptomatic plants (Sieber, 2007; Trivedi *et al*., 2020). The mechanisms by which endophytes transition from commensalism or mutualistic interactions to becoming pathogens remain poorly understood (Douanla-Meli *et al*., 2013; Hardoim *et al*., 2015).

The host microbiome can undergo substantial changes in community structure in the presence of pathogenic species and disease progression (Douanla-Meli *et al*., 2013; Mina *et al*., 2020). For instance, in olive orchards suffering from anthracnose, lower endophyte diversity is observed with higher disease incidence (Martins *et al*., 2021). In Acute Oak decline, the dominant bacterial species was stimulated by a co-invading beetle with additional effects likely caused by other micro-organisms associated with the host or the beetle (Doonan *et al*., 2020). Synergism among different pathogens can increase disease severity in various tree species including apple, chestnut, hazelnut and grapevine (Lamichhane and Venturi, 2015). The impact on tree health resulting from microbial interactions with sequential or cumulative effects can also be modulated by abiotic factors. The deterioration of trees frequently includes abiotic predisposing elements, such as soil microclimate attributes that interact with microbial or insect-induced harm (Doonan *et al*., 2020). Climate factors such as rising temperatures, changes in precipitation patterns, and extreme weather events like droughts and extreme temperatures can induce stress in trees, weaken their immune systems, and make them more vulnerable to pests and other stressors (Camarero *et al*., 2015; Denman *et al*., 2018). Decline diseases, where both abiotic and biotic interactions contribute significantly to disease development, need to be addressed with an integrated system approach (Denman *et al*., 2018).

Grapevine wood dieback is considered the main threat to sustainable grapevine cultivation worldwide but is caused by yet unresolved factors (Cobos *et al*., 2022). A range of wood-colonizing fungal pathogens were suggested to contribute to disease progression (Larignon and Dubos, 1997; Mugnai *et al*., 1999; Larignon *et al*., 2009; Bertsch *et al*., 2013) in addition to changes in climatic and soil conditions (Marchi *et al*., 2006; Surico *et al*., 2006). The main form of dieback is identified as grapevine trunk disease (GTD) with significant impacts on yield and reduced fruit quality leading to high plant replacement rates and economic losses (Bertsch *et al*., 2013; Gramaje *et al*., 2018). GTD is classified into several disease types including the most damaging esca (Bortolami *et al*., 2021). Similar plant declines were also observed on many other woody species including lemon, olive, apple, pomegranate trees without clear associations of potential pathogens and the onset of symptoms (Markakis *et al*., 2017). Esca includes trunk necrosis development in mature vine, along with foliar symptoms and/or symptoms on the shoots with grape wilting. The expression of foliar symptoms can be discontinuous, but plants usually die within a few years following the onset of initial symptoms (Bruez *et al*., 2013; Kenfaoui *et al*., 2022). The discontinuous expression of the disease suggests complex interactions with potential pathogenic species and environmental conditions (Andolfi *et al*., 2011). Esca disease is thought to be associated with the activity of three distantly related fungi (Surico, 2009): *Phaeoacremonium* spp., *Phaeomoniella chlamydospora*, and *Fomitiporia mediterranea*, considered as the most serious pathogens of vines and are the main agents of vascular disease in Europe (Andolfi *et al*., 2011; Bertsch *et al*., 2013; Brown *et al*., 2020). Members of the Botryosphaeriaceae family are also considered to play a role in the disease complex (Bertsch *et al*., 2013; Gramaje *et al*., 2018). These fungi have consistently been isolated from symptomatic grapevines, displaying a close association with esca symptoms such as foliar necrosis and wood discoloration (Mugnai *et al*., 1999; Bruno and Sparapano, 2006). However, fungal species isolated from symptomatic plants often occur both on symptomatic and asymptomatic plants suggesting that the disease is not solely triggered by the presence of specific species (Hofstetter *et al*., 2012; Bruez *et al*., 2016; Del Frari *et al*., 2019). Shared occurrence of fungal species in both symptomatic and asymptomatic plants suggests a potential endophytic phase (Gramaje *et al*., 2018). The exact mechanisms and interactions between these fungi and the grapevine host remain poorly understood (Bertsch *et al*., 2013). Whether the association of fungal species with symptom development of esca is based on causal relationships remains unknown (Bertsch *et al*., 2013; Fischer and Peighami-Ashnaei, 2019). The major limitation of the system is that disease symptoms cannot be reproduced in controlled infections (Reis *et al*., 2016).

A systematic investigation into esca symptom development using standardized grapevine genotypes planted in diverse environments revealed that soil water holding capacity is likely a factor favoring symptom development (Monod *et al*., 2023). Further studies have shown that water availability can influence symptom expression (Marchi *et al*., 2006; Sosnowski *et al*., 2007) and symptom development was likely favored by stronger amplitudes in wet/dry climate transitions (Monod *et al*., 2023). High-throughput amplicon sequencing techniques can generate high-resolution assessments of fungal diversity within grapevine trunks (Dissanayake *et al*., 2018; Pacifico *et al*., 2019; Monod *et al*., 2022). By examining changes in microbiome composition as a function of symptom development in various environments, helps pinpoint species potentially involved in the disease.

Here, we aimed to test the hypothesis that grapevine esca disease progression is trackable by reproducible driver species among fungal communities inhabiting trunks. To achieve this, we analyzed a replicated set of 21 vineyards, all planted simultaneously with a single susceptible cultivar (*i.e.* Gamaret), at the landscape scale. We sampled asymptomatic and symptomatic plants on each vineyard in two different years to perform amplicon sequencing analyses of the grapevine trunk fungal communities. We analyzed mycobiome composition partition within and across vineyards to assess community stability in the absence of disease. Using repeated assessment of asymptomatic and symptomatic plants detected in vineyards, we aimed to investigate potential associations of specific taxa with disease symptom expression.

## Results

### Replicated assessment of the trunk mycobiome across vineyards

We analyzed 21 vineyard plots planted simultaneously in 2003 with *Vitis vinifera* cv. Gamaret in Western Switzerland (Figure 1A). All plants originate from a single nursery to ensure standardization of both age and genetic makeup. The set of replicated Gamaret plots was tracked based on physiological indicators such as yield, must and leaf chemical composition, plant water status, along with meteorological and climatic recordings, soil analyses, and the incidence of esca (Monod *et al*., 2023). Over a span of four years (2018-2021), mortality due to esca was recorded at each site at the onset of plant symptoms and collapse. The mortality rate was highly variable between sites, ranging from 0-47% in 2017 (Figure 1C). To determine whether the grapevine trunk wood fungal community could explain the prevalence of esca, we sampled vine plants in the plot network in 2019 and 2021 (Figure 1B). We randomly selected 10 asymptomatic and 5 symptomatic plants showing either foliar symptoms or apoplexy (Figure 1B, 1D). We used an optimized protocol (Hofstetter *et al*., 2012) to obtain wood cores at the grafting point for each plant (Monod *et al*., 2022). To barcode the endophytic fungal community present in the wood cores, we amplified the ITS with primer pairs ITS1F-ITS4. Previous work on the mycobiome of grapevine trunks showed that utilizing the ITS fragments alone offered a better trade-off between depth of coverage and taxonomic resolution compared to analyzing a longer fragment including also segments of the 28S ribosomal gene sequence (Monod *et al*., 2022).

**Figure 1.**
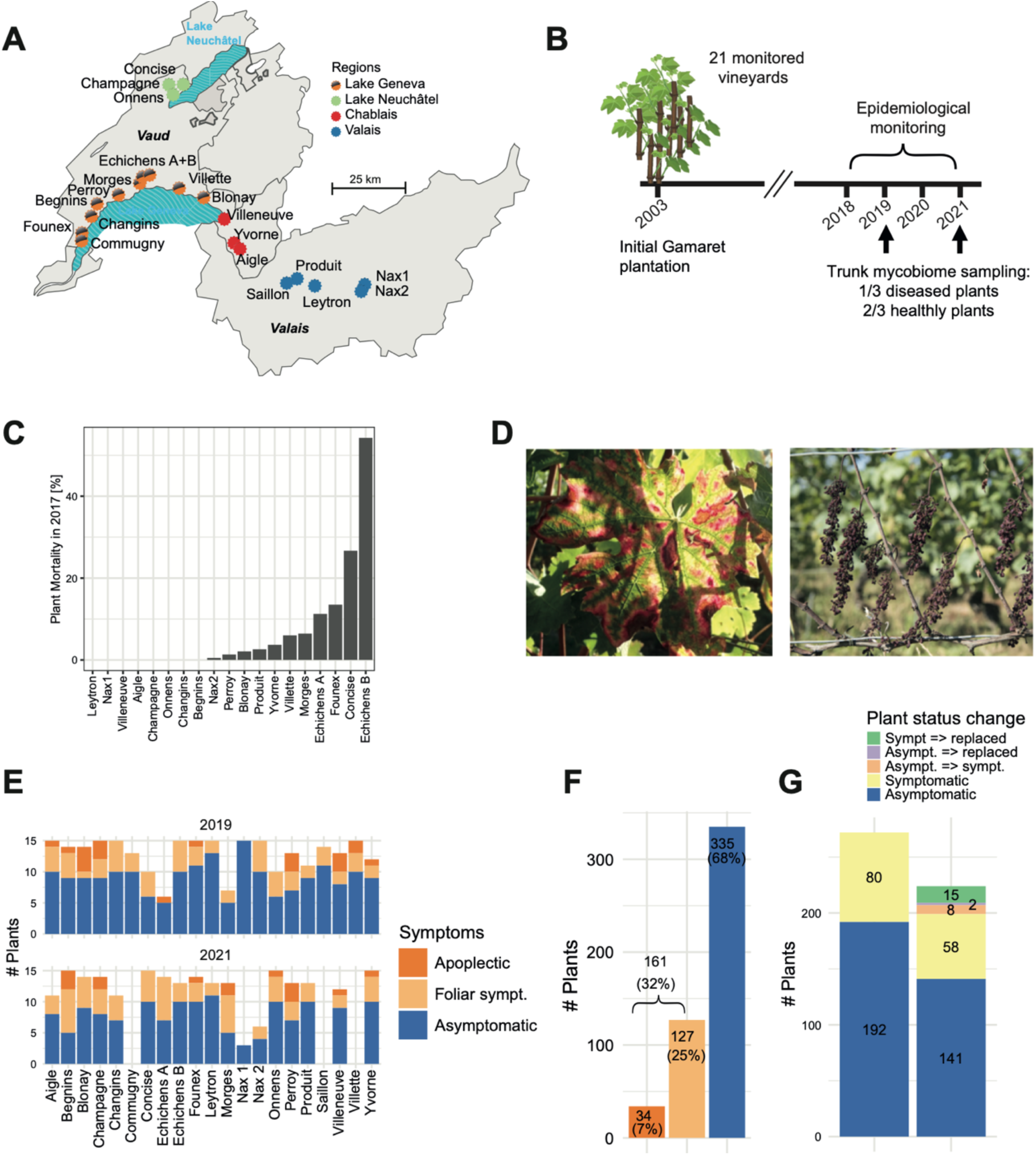
Collection of vine trunk samples to survey the mycobiome community composition. **A)** Location of the studied vineyards (n=21) in Western Switzerland, coloured according to the main viticultural regions. **B)** Life history of the studied vineyards with planting in 2003, followed by the monitoring of esca-BD symptoms (2018-2021) and the sampling seasons (2019 and 2021). **C)** Mortality rates attributed to esca in the studied vineyards (status 2017). **D)** Esca symptoms: typical foliar symptoms (“tiger-stripes”) and apoplectic symptoms with wilting of the whole plant. **E)** Number of samples successfully sequenced according to categories. **F)** Proportion of plants sampled by category (asymptomatic plants n=335, symptomatic plants with leaf symptoms n=127, and symptomatic plants with apoplexy n=34). **G)** Overview of sequenced plants by symptom category and sampling year. The temporal sequence of the health status refers to the observed status in 2019 and 2021.

We successfully amplified 496 samples over all sites (Supplementary Table S1). Samples with low PCR yield were excluded as well as samples from sites where the vineyard was uprooted during the sampling period (see Methods for details). We generated PacBio circular consensus sequencing (CCS) data for 192 asymptomatic and 80 symptomatic plants for the 2019 sampling period (Figure 1G). In 2021, 141 plants (out of 192) had remained asymptomatic and 58 (out of 80) had remained symptomatic. Additionally, we observed that eight plants initially asymptomatic in 2019 became symptomatic (Figure 1G). Furthermore, the plot owners uprooted two asymptomatic and 15 symptomatic plants from 2019. In 2021, we randomly selected new plants as replacement. The composition of the sampled plants, including proportions of asymptomatic and symptomatic individuals varies among plots in particular for esca symptom categories (Figure 1E). Overall, we sequenced 335 asymptomatic plants (68%) and 161 symptomatic plants (32%) (Figure 1F).

We analyzed 3,390,060 CCS reads after quality filtering steps with a mean of 6,834 reads per sample. The reads clustered into 4,129 distinct amplicon sequence variants (ASVs), assigned to 697 species based on matches in the UNITE fungal ribosomal DNA database (Kõljalg *et al*., 2019). The median number of reads per ASV was 25 (Figure 2A). Most AVSs were rare with 30% of the ASVs having 10 reads or less. We obtained a median of 121 ASVs per sample (Figure 2C). The number of detected ASVs per sample was correlated with the number of reads (*r*=0.47, *p*<0.05; Figure 2B; Supplementary Table S2). Overall, the recovered endophytic community was composed of Ascomycota (87.8%), Basidiomycota *(6.82%),* Chytridiomycota (3.29%) and others (2.1%). Among the 41 identified classes, the most abundant classes were Eurotiomycetes (34.4%), Dothideomycetes (30.3%), Sordariomycetes (12.2%), Lecanoromycetes (4.45%) and Leotiomycetes.(4.21%). Among the 497 detected genera, we found a dominance of *Phaeomoniella* (27.6%), *Aspergillus* (5.55%), *Phaeoacremonium* (4.50%), *Pseudopithomyces* (4.01%), *Angustimassarina* (3.90%) (Figure 2D, E, F; Supplementary Table S3).

**Figure 2.**
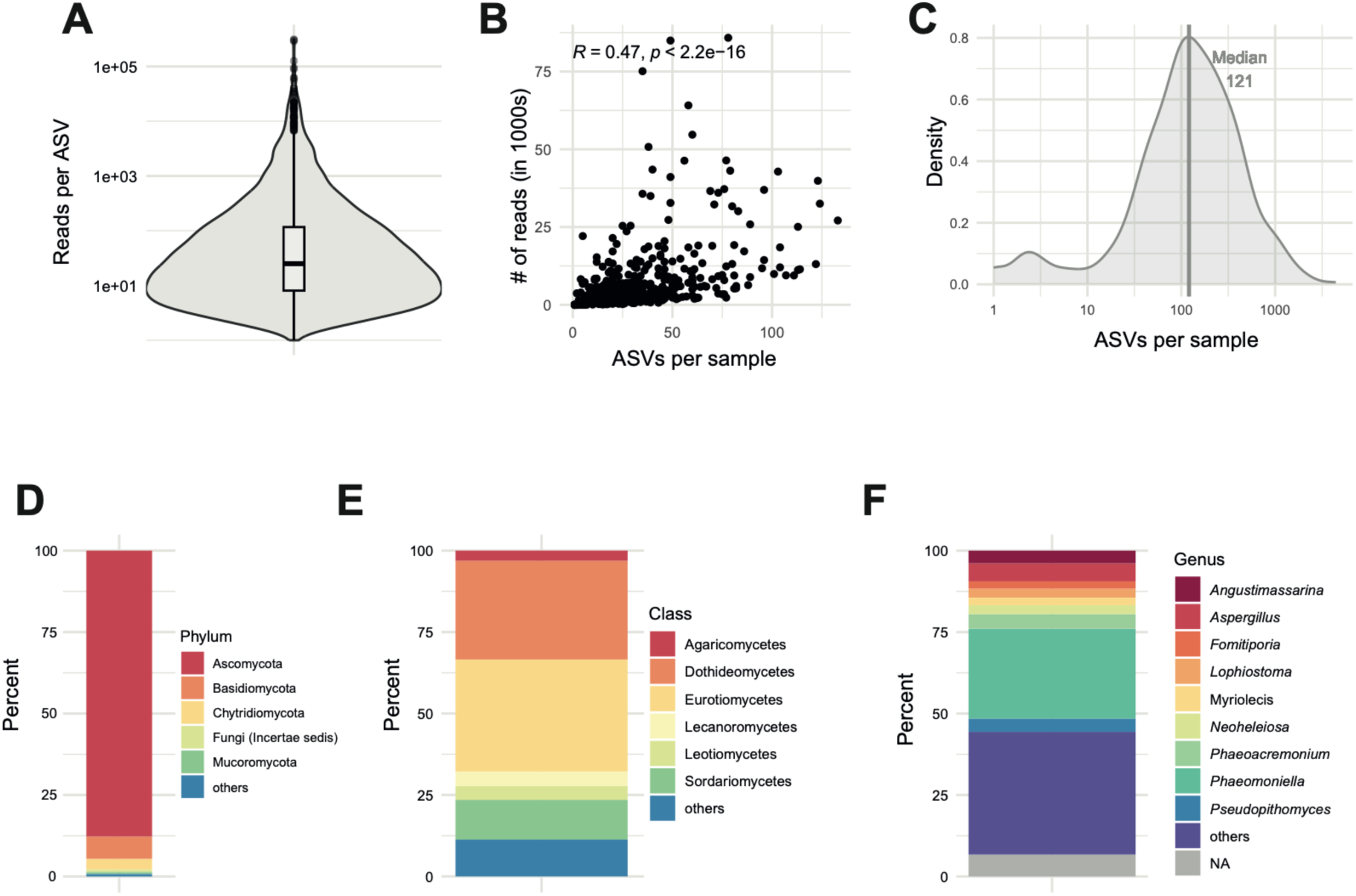
Amplified sequence variants (ASV) recovery and diversity. **A)** Recovered reads per ASV (median= 25 reads)**. B)** Relationship between the number of raw reads and the ASVs detected per sample. **C)** Distribution of ASVs per sample (median=121 ASVs). **D)** Proportion of phyla represented by the amplicon sequences (“others”: phyla with <0.5%). **E)** Proportion of classes in the total sequences (“others“: classes with <2%). **F)** Proportion of genus in the total sequences (“others”: comprises classes with <2% frequency).

### Fungal microbiome structure among healthy and symptomatic plants

Plants are typically associated with diverse microbiomes independent of their health status. To assess the fungal microbiome structure of asymptomatic plants, we analyzed the 502 fungal species detected in asymptomatic grapevine trunks. The diversity of the recovered mycobiome varied between the two sampling years with 1-133 ASVs recovered per plant and a total of 3124 ASVs. The total diversity between the two sampling years was comparable with 351 species recovered in 2019 (1751 ASVs; *n* =192 samples) and 374 species in 2021 (2057 ASVs; n=143 samples. The mycobiome was only weakly shared between regions across Western Switzerland with 158 (3.8%) out of 4129 ASVs found in all regions (Figure 3A). If we consider the proportions of reads associated with each ASVs, fungal communities are more similar with 64% of reads assigned to the same taxa across geographical regions. Differences in mycobiome composition between regions were mostly due to rare ASVs. *P. chlamydospora* was the most abundant species in all regions. A principal coordinate analysis (pCoA) of the mycobiome revealed substantial overlaps among regions and vineyards, yet fungal communities differ significantly among the regions (PERMANOVA, *R*^2^=0.016, *p*=0.001) and among vineyards (PERMANOVA, R2= 0.096, P= 0.001). The pCoA highlights the substantial mycobiome variability among plants, vineyards and regions and the challenge to test for consistent species occurrences across fungal communities (Figure 3B).

**Figure 3.**
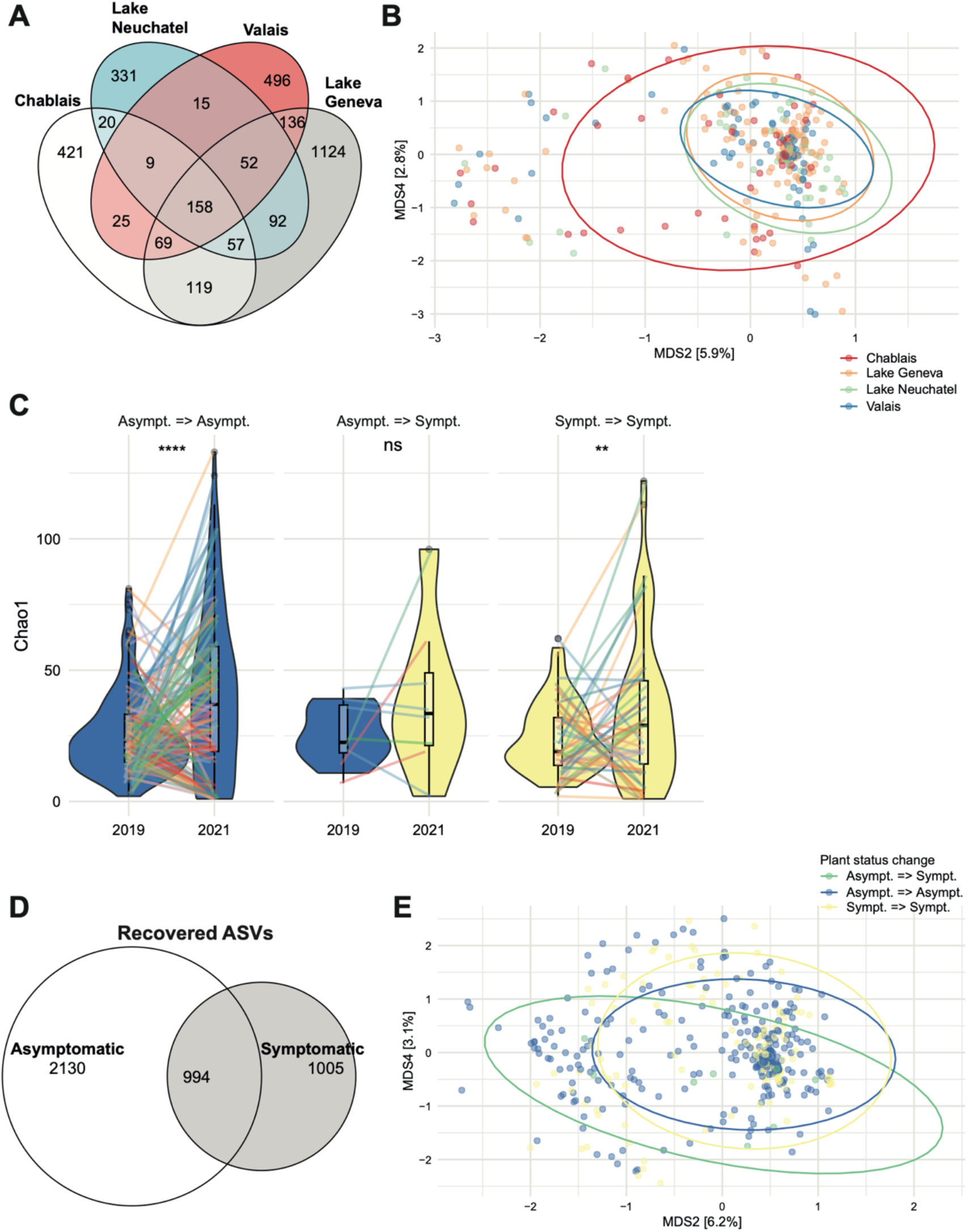
Diversity of asymptomatic or symptomatic plant mycobiomes. **A)** Proportion of ASVs shared among asymptomatic plants per geographic regions. **B)** Principal coordinate analysis (PCoA, no transformation, Bray-Curtis distance on ASV diversity, *n* = 3124) of mycobiome diversity of asymptomatic plants across geographic regions. Each point represents the mycobiome composition at the ASV level of the trunk of once sampled vine plants. **C)** Violin plots displaying the α-diversity on esca asymptomatic and symptomatic sampled plants (Chao1 index) taking into account their epidemiological history (Asympt => Asympt: plants that remained asymptomatic during the two years of sampling; Asympt => Sympt: plants that changed from asymptomatic in 2019 to symptomatic in 2021; Sympt_Sympt: plants recorded as symptomatic during the two years of sampling) with individual samples linked and colour-marked by vineyard. **D)** Proportion of ASVs shared between asymptomatic and symptomatic sampled plants. **E)** Principal coordinate analysis (PCoA, no transformation, Bray-Curtis distance on ASVs diversity, *n* = 4169) of the mycobiome diversity of the samples.

Symptomatic plants (foliar and apoplexy symptoms) did not differ significantly from asymptomatic plants in recovered species or ASV diversity with 502 species (3124 ASVs) detected among asymptomatic plants (*n*=335) and 418 species (1999 ASVs) detected among symptomatic plants (*n*=161) (ANOVA, *p*>0.05). Comparisons of Chao1 diversity among different plant health status categories revealed significant differences between plants remaining healthy (*i.e.* asymptomatic) or keep showing symptoms between the sampling years (Wilcoxon *p*=1.6e^-5^ for asymptomatic plants; *p=*0.061 for symptomatic plants; Figure 3C). Variability in the recovered diversity is not associated with the health status of the sampled plant. The fungal diversity recovered for the same plant varied across the two time points but differences in diversity for plants turning from asymptomatic to symptomatic across sampling years were not significant (Chao1 diversity index; Wilcoxon p>0.5; Figure 3C). However, this assessment is based on a comparatively low number of observations (*n* = 8). Asymptomatic and symptomatic plants shared overall 24% of ASVs (Figure 3D). If we consider the proportions of reads associated with each ASVs, fungal communities are more similar with 89% of reads assigned to the same taxa between health status. Differences in fungal community composition across samples of different health status were largely due to rare taxa. The pCoA revealed no obvious clustering between plants remaining asymptomatic, persistant in a symptomatic stage or turning symptomatic over the sampling period (Figure 3E). Community composition was nevertheless significantly different between symptomatic and asymptomatic plants (PERMANOVA, *R*^2^=0.003, *p*=0.014). Fungal community composition was significantly different between asymptomatic plants compared to plants suffering from apoplexy (PERMANOVA, *R^2^*=0.00465, *p*=0.005). Fungal communities of plants presenting either foliar symptoms or apoplexy were not significantly different (PERMANOVA, p>0.05).

### Taxonomic profiles highlight taxa linked to asymptomatic plants

We assessed evidence of significantly overrepresented taxa in either asymptomatic and symptomatic plants using discriminant analyses (DA; Supplementary Table S4). DA indices were constructed for a total of 496 plant samples. We used three different approaches to assess evidence for taxa enrichment in symptomatic versus asymptomatic plant trunk mycobiome. First, linear discriminant analysis effect size (LEfSe) analyses identified the *Neosetophoma* genus, a species of the same genus (*N. shoemaker*) and the related Phaeosphaeriaceae family and the class of Tremellomycetes as enriched in asymptomatic plants (Figure 4A). Second, an analysis of composition of microbiomes (ANCOM) identified six enriched genera including two enriched in asymptomatic plants (*Neosetophoma* and *Filobasidium*) and three enriched in symptomatic plants (*Tausonia*, *Verrucoccum* and *Mortierella*). ANCOM also identified the Asycomycota phylum as enriched in symptomatic samples (Figure 4B). ANCOM-BC identified seven genera (*Neosetophoma*, *Calloriaceae*, *Naganishia*, *Curvibasidium*, *Trichoderma*, *Cyphellophora* and *Lophiostoma*) and an order (Pleosporales) as having reduced abundance (negative LFC) in symptomatic compared to asymptomatic plants (Figure 4C). Hence, the ANCOM method was the only one to identify enriched taxa in symptomatic plants. The *Neosetophoma* genus was supported by evidence and consensus between methods for association with asymptomatic plants. Proportionally, *Neosetophoma* genus represents 0.5% of the reads of asymptomatic plants compared to 0.03% in symptomatic plants (Figure 5C). Upon examining the distribution of the *Neosetophoma* genus across various geographical regions, a notably higher occurrence was observed in the Valais region (Figure 5D). Valais is the region that exhibits the lowest incidence of esca impact (Monod *et al*., 2023). When we examined the presence of the *Neosetophoma* genus alongside the recorded mortality rates in each vineyard, we did not detect any correlation though (R=0.05 p=0.518) (Figure 5E).

**Figure 4.**
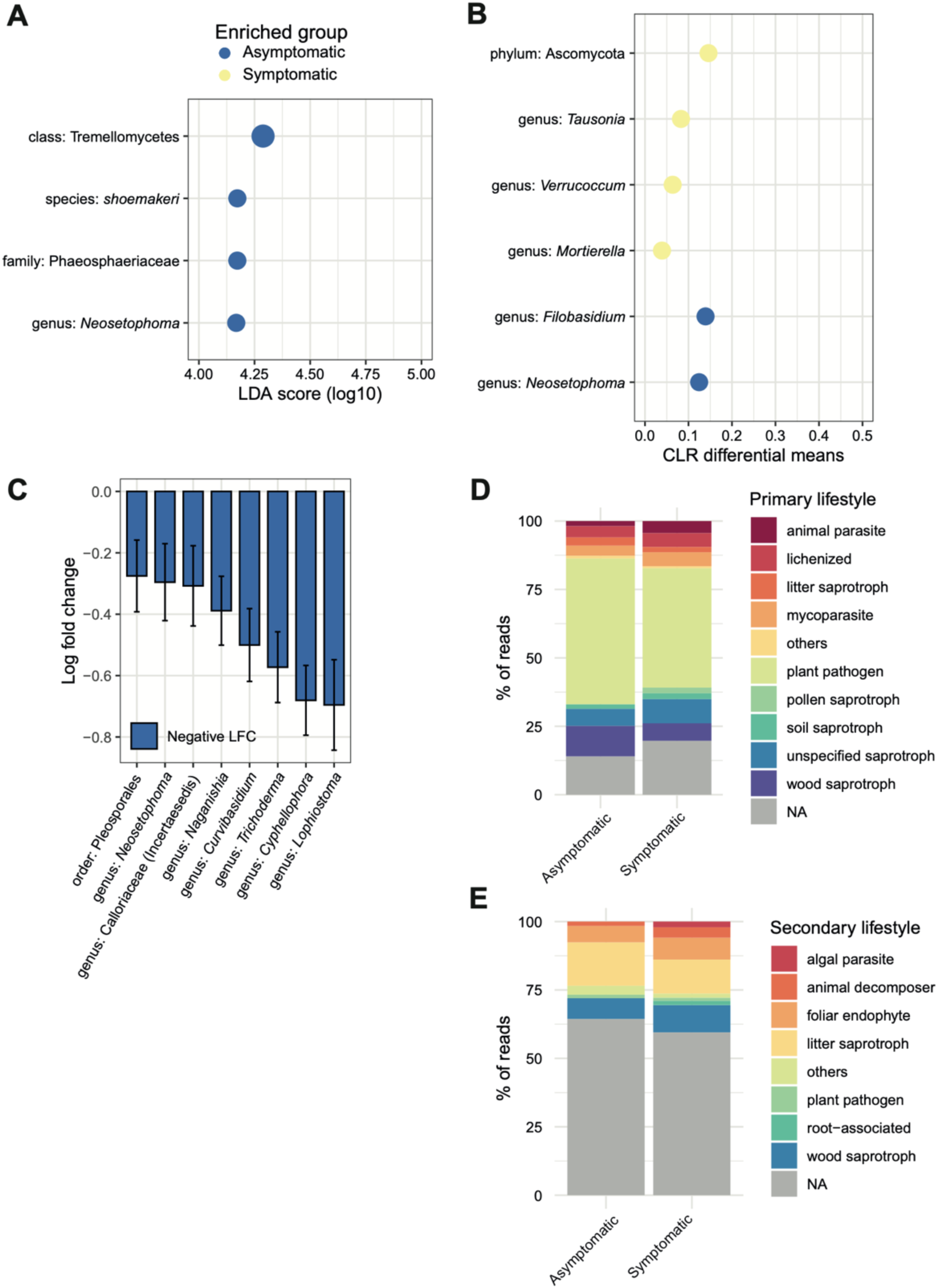
Identification of differentially abundant taxa in the mycobiome of asymptomatic and symptomatic plants. **A)** Linear discriminant analysis (LEfSe) was used to identify overabundant taxa in asymptomatic and symptomatic plants (CLR normalization, p-value < 0.05). Enriched taxa in the asymptomatic group are shown in blue, enriched taxa in the symptomatic group in yellow. The list of discriminating features according to the classes (asymptomatic and symptomatic) is ordered by the magnitude of the effect with which they differentiate the classes. **B)** Microbiome composition analysis (ANCOM) identified compositional differences (p-value < 0.05) in the mycobiome communities of asymptomatic and symptomatic sampled plants. Enriched taxa in the asymptomatic group are shown in blue, enriched taxa in the symptomatic group in yellow. **C)** Analysis of microbiome composition represented by effect size (log fold change) and 95% confidence interval bars (two-sided; Bonferroni adjusted) derived from the ANCOM-BC model. **D)** Traits based approach with proportion of primary lifestyle between asymptomatic and symptomatic sampled plants. **E)** Traits based approach with proportion of secondary lifestyle between asymptomatic and symptomatic sampled plants.

**Figure 5.**
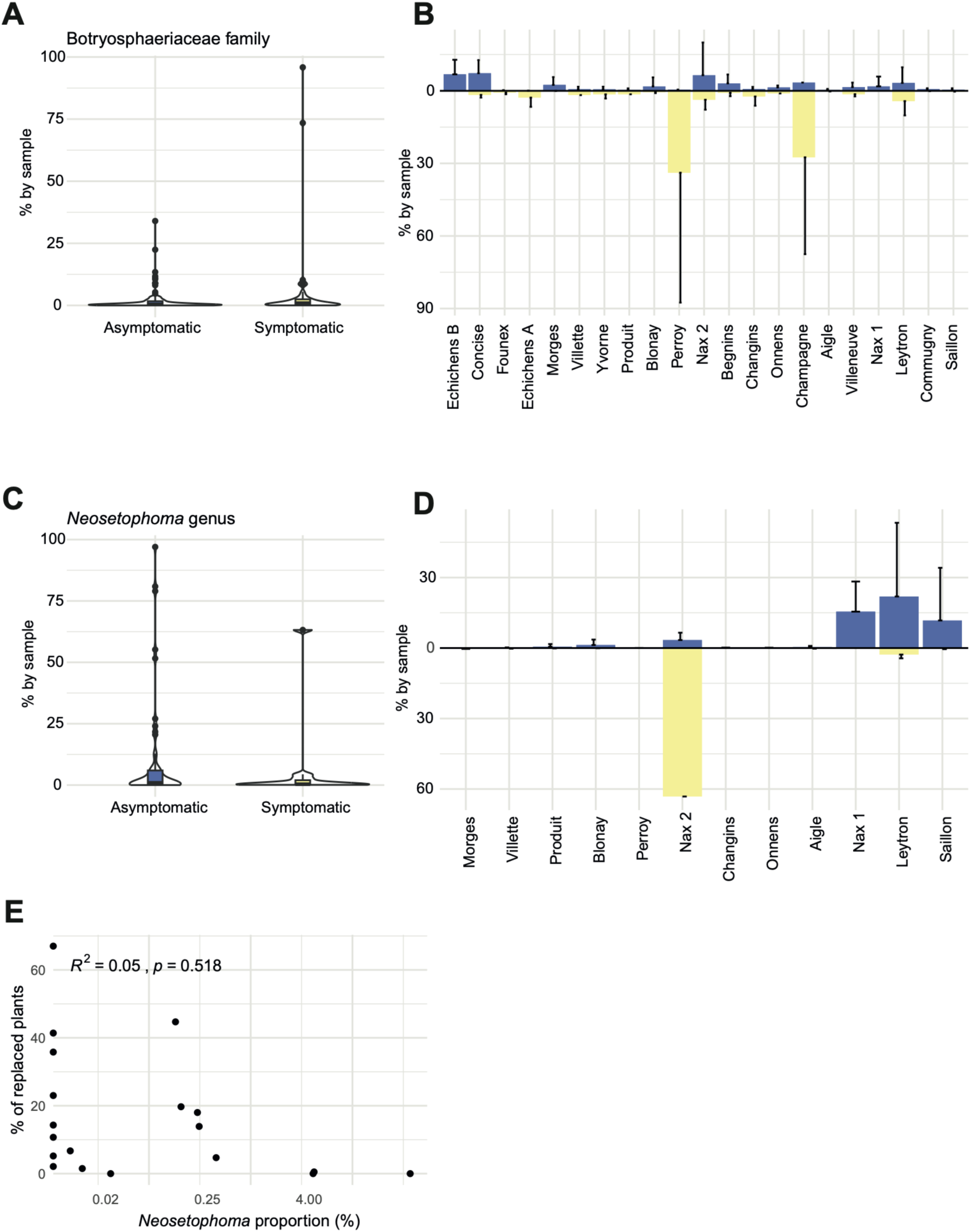
A) Botryosphaeriaceae family with symptomatic and asymptomatic taxa. **B)** The mean (standard deviation in black) of the Botryosphaeriaceae family proportion by vineyard for asymptomatic (blue) and symptomatic (yellow) sampled plants. **C)** Proportion of *Neosetophoma* genus between asymptomatic and symptomatic plants. **D)** The proportion of the *Neosetophoma* genus varies between vineyards in asymptomatic (blue) and symptomatic (yellow) sampled plants. **E)** Relationship between the proportion of *Neosetophoma* genus by plot and the proportion of replaced plants.

To determine potential differences in the ecological roles of fungal communities in symptomatic and asymptomatic plant samples, we classified each identified genus into functional groups using the FungalTrait database (Põlme *et al*., 2020). After considering predicted guilds and trophic modes, our analysis revealed no significant differentiation between asymptomatic and symptomatic plants (Figure 4D, E). It should be noted that many genera are classified as pathogenic. This is potentially a bias of the database, as pathogenic genera are more studied and described than others.

### Key genera previously associated with esca

We retrieved a set of taxa commonly described to be associated with grapevine trunk diseases (Andolfi *et al*., 2011; Bertsch *et al*., 2013; Brown *et al*., 2020). We focused on presence and relative abundance of the genera *Phaeomoniella*, *Phaeoacremonium* and *Fomitiporia*, as well as the Botryosphaeriaceae family (Figure 5A-B; Figure 6). We retrieved ASVs assigned to each taxonomic unit across vineyards and plant health status. We found no evidence that these taxonomic units were enriched in symptomatic plants (Figure 6). The proportions of *Phaeomoniella* (mean of 34.1% in asymptomatic; 33.1% in symptomatic plants), *Phaeoacremonium* (mean of 14.7% in asymptomatic; 11.7% in symptomatic plants) and *Fomitiporia* (mean of 16.5% in asymptomatic; 11.6% in symptomatic plants) in both symptomatic and asymptomatic plants were comparable (Figure 6 A, C, E). Relative abundance of the focal taxa varied across vineyards ranging for *Phaeomoniella* from 100% to 0.02% in asymptomatic and symptomatic plants, for *Phaeoacremonium* from 100% to 0.02% in asymptomatic plants and from 100% to 0.06% in symptomatic plants and for *Fomitiporia* from 95.6% to 0.03% in asymptomatic plants and 77,5% to 0.02% in symptomatic plants (Figure 6 B, D, F).

**Figure 6.**
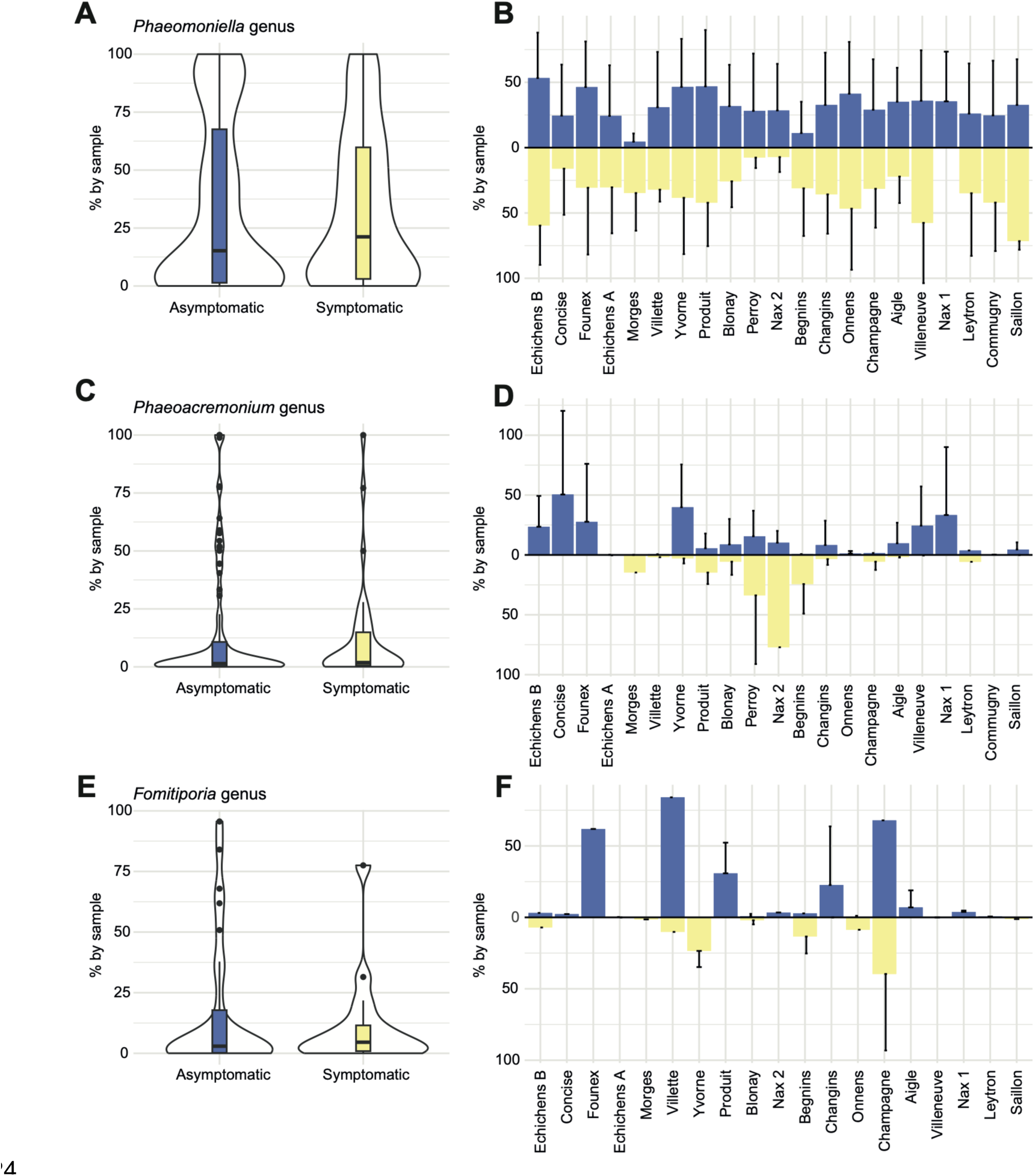
Fungal taxa commonly associated with esca per genus. **A)** *Phaeomoniella* genus with symptomatic and asymptomatic taxa. **B)** The mean (standard deviation in black) of the *Phaeomoniella* genus proportion by vineyard for asymptomatic (blue) and symptomatic (yellow) sampled plants. **C)** Proportion of *Phaeoacremonium* genus between asymptomatic and symptomatic plants. **D)** The proportion of the *Phaeoacremonium* genus varies between vineyards in asymptomatic (blue) and symptomatic (yellow) sampled plants. **E)** Proportion of the *Fomitiporia* genus between asymptomatic and symptomatic sampled plants. **F)** The proportion of the *Fomitiporia* genus across vineyards and asymptomatic (blue) and symptomatic (yellow) sampled plants.

## Discussion

Woody plant decline is likely caused by a multitude of fungal species and is facilitated by environmental conditions. Here, our objective was to examine compositional differences in trunk-inhabiting fungal communities in vineyards affected to varying degrees by esca. The analysis of the vine trunk mycobiome revealed a remarkably diverse fungal community with weak differentiation at the vineyard of regional level. We found overrepresentation of several taxa in asymptomatic plants; however no taxa were overrepresented in symptomatic plants. Additionally, key taxa typically implicated in esca did not show any significant association with plant health status.

### Extensive mycobiome variability across sampling scales

Fungal diversity among sampled plants exhibited high heterogeneity confirming analyses conducted across various wild or cultivated plant species (Vandenkoornhuyse *et al*., 2015; Pozo *et al*., 2021). The grapevine trunk mycobiome primarily consisted of rare taxa, consistent with many host-associated mycobiome studies (Gobet *et al*., 2010; Segata *et al*., 2011; Lundberg *et al*., 2012; Travadon *et al*., 2013; Vaz *et al*., 2018; Del Frari *et al*., 2019). Variation in sample diversity may be attributed to factors such as sampling bias, disparities between plant tissues containing both living and deceased material, intra-vineyard diversity, or differences in pedoclimatic conditions (Vandenkoornhuyse *et al*., 2015; Pacifico *et al*., 2019; Bettenfeld *et al*., 2022). We observed dissimilarities in fungal composition across geographic areas consistent with findings from previous studies on grapevine microbiome composition (Bekris *et al*., 2021) and other woody plants (Proença *et al*., 2017). These observations suggest that fungal endophytes colonize the tissues of the hosts through a potential horizontal transfer of diversity from the surrounding environment via soil- or airborne spores (Saikkonen *et al*., 2004; Vaz *et al*., 2018; Rana *et al*., 2019).

Alpha diversity declined with increasing disease symptom severity. Reduced diversity was shown in other systems to play a protective role mediated by the remaining species (Koskella *et al*., 2017). Similar findings were obtained from acute oak decline (Doonan *et al*., 2020), from fungal root pathogens (Mendes *et al*., 2011) or bumble bees (Koch and Schmid-Hempel, 2011). The opposite was also observed though in pine wilt disease (Proença *et al*., 2017) or ash dieback (Griffiths *et al*., 2020) where higher diversity of the microbiome was observed along with symptom severity. Higher microbiome diversity is thought to stem from the pathogen suppressing plant resistance mechanisms and, thereby, facilitating the colonization by other microorganisms (Proença *et al*., 2017). Plants affected by esca showed neither a decrease or increase in alpha diversity in our study. No significant differences in alpha diversity of declined and healthy trees were found in key tree species (holm oak, cork oak, chestnut and pyrenean oak) of the Mediterranean forest (Diez-Hermano *et al*., 2022). While richness in diversity remained unchanged, alterations in community composition could still have occurred depending on the health status of the plant. However, the interpretation of the functional role of the fungal community can be challenging because fungal species can undergo lifestyle transitions depending on the environment (Sieber, 2007; Romeralo *et al*., 2022).

### Grapevine fungal community structures are shaped by rare taxa

Despite no overall diversity effects, we detected a broad range of alterations in the community composition between asymptomatic and symptomatic plants. However, plant health was not a strong enough factor to reveal distinct community effects using beta dissimilarity analyses. Across geography and esca health status, we observed high inter-sample variability. Such variability in the host-associated mycobiome creates significant challenges to pinpoint cryptic species underpinning diseases. Nonetheless, many environmental microbiome communities are typically characterized by the presence of a long tail of rare taxa (Pedrós-Alió, 2006; Gobet *et al*., 2010). Overcoming statistical limitations in associating rare taxa with disease development would require either substantially expanding the sampling effort or reducing environmental noise.

### No differentiated fungal community associated with symptomatic plants

We conducted analyses to assess the enrichment of particular taxonomic groups in symptomatic plants using three distinct approaches but found no strongly associated taxa. This is in line with previous research on fungal trunk communities affected by esca, revealing no direct association between specific taxa and symptomatic esca plants (Hofstetter *et al*., 2012; Bruez *et al*., 2014; Del Frari *et al*., 2019). Our study builds upon previous research by substantially increasing the number of samples, emphasizing that even sampling strategies with hundreds of data points may still be underpowered. Previous research conducted on the same vineyards has linked the incidence of esca symptoms to pedo-climatic factors (Monod *et al*., 2023), suggesting that soil water holding capacity is a key factor for disease development. Soil retention capacity is influenced by the amount of precipitation and various soil characteristics. Whether such soil properties are causal or merely show correlated responses to an, as yet, unknown factor remains uncertain. If soil properties are indeed the root cause of the disease, a number of fungal taxa may in turn sporadically associate with particular soil types without playing a relevant role in the disease. Furthermore, any association of endophyte taxa may similarly be due to correlations between soil characteristics and fungal diversity (Geiger *et al*., 2022). Furthermore, endophytes may transition from a latent asymptomatic state to an active state after the plant encountered stress conditions such as drought, humidity, or nutrient starvation (Mishra *et al*., 2021). An interesting observation was the significant enrichment of the *Neosetophoma* genus in asymptomatic plants, as supported by consensus among all differential abundance methods. This genus is most prevalent in the Valais region, which also exhibits the lowest incidence of esca. However, it is worth noting that the *Neosetophoma* genus is absent from numerous analyzed vineyards. Therefore, the strong relationship between the presence of the *Neosetophoma* genus and the absence of esca symptoms should be interpreted cautiously. It is conceivable that endophytes residing in woody plants play a defensive role for the host plant by producing a range of protective mycotoxins and enzymes (Pacifico *et al*., 2019; Stewart *et al*., 2021). Lack of associated taxa with trunk disease does not negate causal interactions of fungal taxa with the disease but rather suggests that the complexity of biotic drivers is too high.

Plant health or disease should not be viewed as a binary concept but rather an expression of symptoms along a continuum. In complex diseases like tree dieback, multiple factors are likely involved and resolving causal relationships between taxa and health is challenging. The presence of fungal endophytes residing within plants without causing harm, challenges our traditional understanding of plant infection processes and how causal taxa should be identified (Mishra *et al*., 2021). The ecological relevance of rare species is increasingly recognized with key functions in host-associated microbiomes (Jousset *et al*., 2017). Yet, determining perturbations caused by rare species is challenging as most studies were likely underpowered (Säterberg *et al*., 2019). Further research under more controlled conditions is needed to determine what disruption or imbalance in the plant microbiome is considered detrimental to plant health (Romani, 2011; Begum *et al*., 2022) and what meaningful boundaries can be drawn between endophytes and pathogens (Lòpez-Fernàndez *et al*., 2015).

## Methods

### Sample collection

Wood samples were collected in August 2019 and 2021 from vineyards located in four viticultural regions in western Switzerland. A total of 21 vineyards planted with the Gamaret variety were sampled. Gamaret originates from a cross between Gamay and Reichensteiner varieties grafted onto 3309C rootstock (*V. riparia* X *V. rupestris*). All plants originate from the same nursery (Les frères Dutruy SA; Founex, Switzerland) and were planted in 2003. The 21 vineyards had been under similar viticultural management. In each vineyard, five symptomatic plants displaying the typical foliar esca symptoms including leaf discoloration, tiger-stripe pattern and plant wilting (Mugnai *et al*., 1999) were collected alongside ten asymptomatic plants in 2019 and 2021. Plants were 16 and 18 years old when samples were collected in 2019 and 2021, respectively. Some vineyards were uprooted (*i.e.,* Villette and Saillon vineyards) and some replacements were managed inconsistently (*i.e.,* Commugny), hence three vineyards were excluded for the second sampling year. Each vine plant selected was sampled at the grafting point using a nondestructive method (Hofstetter et al 2012). A 0.5 cm^2^ piece of bark was removed with a surface-sterilized scalpel (80% ethyl alcohol). Next, sampling was performed using a power drill with a surface-sterilized drill bit (Ø 3.5 mm) at the spot where the bark was removed. Coiled wood (∼60 mg) extracted by the power drill was collected in Eppendorf tubes held underneath using sterilized tweezers. Eppendorf tubes containing the coiled wood were stored at -80°C.

### DNA extraction from wood samples

Eppendorf tubes containing wood samples and two 5-mm iron beads were placed in liquid nitrogen. Material was ruptured two times for 1 min at 30 Hz in a TissueLyser (Qiagen Inc., Germantown, MD, USA). Between and after these two steps of tissue disruption, tubes were placed in liquid nitrogen for 1 min. The tubes were placed on ice for slow thawing and 1 mL of cetyltrimethylammonium bromide (CTAB) was added to each tube. The samples were then centrifuged for 1 min at 15,000 rpm and the supernatant was transferred to a new tube. Fungal DNA was extracted using a Qiacube robot with the DNeasy Plant Pro Kit 69206 (Qiagen).

*Amplification of fungal ribosomal DNA*.

The ITS was targeted for amplification using primers ITS1F (CTTGGTCATTTAGAGGAAGTAA) and ITS4 (TCCTCCGCTTATTGATATGC) (Gardes and Bruns, 1993). We followed the PacBio procedure using barcoded universal primers for multiplexing amplicons, which includes two PCR steps (see https://www.pacb.com). The first PCR program was 30 s of denaturation at 98°C and then 30 cycles of 15 s at 98°C, 15 s at 55°C, and 1 min 30 s at 72°C, followed by a final elongation step for 7 min at 72°C. The second PCR program was 30 s of denaturation at 98°C and then 20 cycles of 15 s at 98°C, 15 s at 64°C, and 1 min 20 s at 72°C, followed by a final elongation step for 7 min at 72°C (Pacific Biosciences, 2019). The final libraries were quantified with a Qubit fluorometer (Thermo Fisher, Foster City, CA, USA), and then all samples were pooled equimolarly. Amplicons were purified and prepared for SMRT sequencing at the Functional Genomics Center in Zürich (FGCZ), Switzerland. Sequencing was performed on the PacBio Sequel II platform.

### Demultiplexing and analyses of amplicon sequence variants

Raw reads were processed with the DADA2 package in R (Callahan Github, https://github.com/benjjneb/dada2). We quality-trimmed, filtered reads and inferred amplicon sequence variants (ASVs) with DADA2. For chimera detection we observed a too light detection made by DADA2 algorithm (isBimeraDenovo) and consequently we used QIIME2 uchime-denovo function. Taxonomic assignments were performed with the function *AssignTaxonomy()* of the DADA2 pipeline, which classifies sequences based on the reference training data sets and based on the UNITE general FASTA release database (2023-07-25, (Abarenkov *et al*., 2023). We calculated Chao1 indices to assess the richness and diversity of the trunk mycobiome using the *vegan* R package (Oksanen et al 2022). Differences in taxonomic diversity were tested using a Wilcoxon test (*p* < 0.05).

### Analyses and characterization of taxa associated with esca

We assessed dissimilarity distances to visualize beta diversity and quantified differences in the overall ASV composition based on a principal coordinates analysis (PCoA) with Bray-Curtis distances (Bray and Curtis, 1957). Beta diversity dissimilarities in fungal communities were assessed at the sample level, health status and geographic region. Differences between groups in taxonomic composition were tested using three distinct methods. Linear discriminant analysis effect size (LEfSe) (Segata *et al*., 2011) is based on Kruskal-Wallis rank sum tests to identify taxa with significant differential abundance (alpha = 0.05) between groups using one-against-all comparisons. The analyses are followed by a linear discriminant analysis (LDA) to estimate the effect size of each differentially abundant feature (LDA >2). ANCOM2 based on Aitchison’s methodology uses relative abundances to infer absolute abundances (Mandal et al 2015). Analysis of Compositions of Microbiomes with Bias Correction (ANCOM-BC) was used with an adjustment for sampling fraction by adding a sample-specific offset term in a linear regression model (Lin and Peddada, 2020). This offset term corrects for biases and the log-transformed linear regression framework addresses microbiome data compositionality (Lin and Peddada, 2020). Niche characteristics and traits shared by identified genera were analyzed using the FungalTraits database (Põlme *et al*., 2020). Figure panels were generated using the R package *ggplot2* v3.3.3 (Wickham).

## Data availability

All PacBio sequencing data are available from the NCBI Sequence Read Archive (SRA).

## Acknowledgments

Library preparation and sequencing were performed at the Functional Genomics Centre Zurich (FGCZ). We are grateful to the members of the mycology research group of Agroscope (N. Lecoultre, E. Michellod, A.-L. Fabre, P.-H. Dubuis, D. Restori, and A. Melgar) for their assistance in sampling vineyards. We are grateful for discussions with Luzia Stadler and Hanna Glad about analysis methods.

## Author contributions

VM, VZ, OV, KG, VH and DC conceived the study; VM collected the data; VM and DC analyzed the data; VM and DC led the writing of the manuscript. All authors commented on the manuscript.

## References

Abarenkov, K., Zirk, A., Piirmann, T., Pöhönen, R., Ivanov, F., Nilsson, R.H., and Kõljalg, U. (2023) UNITE general FASTA release for Fungi.

Andolfi, A., Mugnai, L., Luque, J., Surico, G., Cimmino, A., and Evidente, A. (2011) Phytotoxins produced by fungi associated with grapevine trunk diseases. Toxins 3: 1569–1605.

Bacon, C.W., Glenn, A.E., and Yates, I.E. (2008) Fusarium verticillioides: managing the endophytic association with maize for reduced fumonisins accumulation. Toxin Rev 27: 411–446.

Begum, N., Harzandi, A., Lee, S., Uhlen, M., Moyes, D.L., and Shoaie, S. (2022) Host-mycobiome metabolic interactions in health and disease. Gut Microbes 14: 2121576.

Bekris, F., Vasileiadis, S., Papadopoulou, E., Samaras, A., Testempasis, S., Gkizi, D., et al. (2021) Grapevine wood microbiome analysis identifies key fungal pathogens and potential interactions with the bacterial community implicated in grapevine trunk disease appearance. Environ Microbiome 16: 23.

Bertsch, C., Ramírez-Suero, M., Magnin-Robert, M., Larignon, P., Chong, J., Abou-Mansour, E., et al. (2013) Grapevine trunk diseases: complex and still poorly understood. Plant Pathol 62: 243– 265.

Bettenfeld, P., Cadena I Canals, J., Jacquens, L., Fernandez, O., Fontaine, F., van Schaik, E., et al. (2022) The microbiota of the grapevine holobiont: A key component of plant health. J Advert Res 40: 1–15.

Bettenfeld, P., Fontaine, F., Trouvelot, S., Fernandez, O., and Courty, P.-E. (2020) Woody Plant Declines. What’s Wrong with the Microbiome? Trends Plant Sci 25: 381–394.

Bortolami, G., Gambetta, G.A., Cassan, C., Dayer, S., Farolfi, E., Ferrer, N., et al. (2021) Grapevines under drought do not express esca leaf symptoms. Proc Natl Acad Sci U S A 118:.

Bray, J.R. and Curtis, J.T. (1957) An Ordination of the Upland Forest Communities of Southern Wisconsin. Ecol Monogr 27: 326–349.

Brown, A.A., Lawrence, D.P., and Baumgartner, K. (2020) Role of basidiomycete fungi in the grapevine trunk disease esca. Plant Pathol 69: 205–220.

Bruez, E., Baumgartner, K., Bastien, S., Travadon, R., Guérin-Dubrana, L., and Rey, P. (2016) Various fungal communities colonise the functional wood tissues of old grapevines externally free from grapevine trunk disease symptoms. Aust J Grape Wine Res 22: 288–295.

Bruez, E., Lecomte, P., Grosman, J., Doublet, B., Bertsch, C., Fontaine, F., et al. (2013) Overview of grapevine trunk diseases in France in the 2000s. Phytopathol Mediterr 52: 262–275.

Bruez, E., Vallance, J., Gerbore, J., Lecomte, P., Da Costa, J.-P., Guerin-Dubrana, L., and Rey, P. (2014) Analyses of the temporal dynamics of fungal communities colonizing the healthy wood tissues of esca leaf-symptomatic and asymptomatic vines. PLoS One 9: e95928.

Bruno, G. and Sparapano, L. (2006) Effects of three esca-associated fungi on Vitis vinifera L.: III. Enzymes produced by the pathogens and their role in fungus-to-plant or in fungus-to-fungus interactions. Physiol Mol Plant Pathol 69: 182–194.

Camarero, J.J., Gazol, A., Sangüesa-Barreda, G., Oliva, J., and Vicente-Serrano, S.M. (2015) To die or not to die: early warnings of tree dieback in response to a severe drought. J Ecol 103: 44–57.

Cobos, R., Ibañez, A., Diez-Galán, A., Calvo-Peña, C., Ghoreshizadeh, S., and Coque, J.J.R. (2022) The Grapevine Microbiome to the Rescue: Implications for the Biocontrol of Trunk Diseases. Plants 11:.

Cohen, W.B., Yang, Z., Stehman, S.V., Schroeder, T.A., Bell, D.M., Masek, J.G., et al. (2016) Forest disturbance across the conterminous United States from 1985–2012: The emerging dominance of forest decline. For Ecol Manage 360: 242–252.

Compant, S., Samad, A., Faist, H., and Sessitsch, A. (2019) A review on the plant microbiome: Ecology, functions, and emerging trends in microbial application. J Advert Res 19: 29–37.

Del Frari, G., Gobbi, A., Aggerbeck, M.R., Oliveira, H., Hansen, L.H., and Ferreira, R.B. (2019) Characterization of the Wood Mycobiome of Vitis vinifera in a Vineyard Affected by Esca. Spatial Distribution of Fungal Communities and Their Putative Relation With Leaf Symptoms. Front Plant Sci 10: 910.

Denman, S., Doonan, J., Ransom-Jones, E., Broberg, M., Plummer, S., Kirk, S., et al. (2018) Microbiome and infectivity studies reveal complex polyspecies tree disease in Acute Oak Decline. ISME J 12: 386–399.

Diez-Hermano, S., Ahmad, F., Niño-Sanchez, J., Benito, A., Hidalgo, E., Escudero, L.M., et al. (2022) Health condition and mycobiome diversity in Mediterranean tree species. Frontiers in Forests and Global Change 5:.

Dissanayake, A.J., Purahong, W., Wubet, T., Hyde, K.D., Zhang, W., Xu, H., et al. (2018) Direct comparison of culture-dependent and culture-independent molecular approaches reveal the diversity of fungal endophytic communities in stems of grapevine (Vitis vinifera). Fungal Divers 1: 85–107.

Doonan, J.M., Broberg, M., Denman, S., and McDonald, J.E. (2020) Host-microbiota-insect interactions drive emergent virulence in a complex tree disease. Proc Biol Sci 287: 20200956.

Douanla-Meli, C., Langer, E., and Mouafo, F.T. (2013) Fungal endophyte diversity and community patterns in healthy and yellowing leaves of Citrus limon. Fungal Ecol.

Fischer, M. and Peighami-Ashnaei, S. (2019) Grapevine, esca complex, and environment: the disease triangle. Phytopathol Mediterr 58: 17–37.

Gardes, M. and Bruns, T.D. (1993) ITS primers with enhanced specificity for basidiomycetes--application to the identification of mycorrhizae and rusts. Mol Ecol 2: 113–118.

Geiger, A., Karácsony, Z., Golen, R., Váczy, K.Z., and Geml, J. (2022) The compositional turnover of grapevine-associated plant pathogenic fungal communities is greater among intraindividual microhabitats and terroirs than among healthy and Esca-diseased plants. Phytopathology 112: 1029–1035.

Gobet, A., Quince, C., and Ramette, A. (2010) Multivariate Cutoff Level Analysis (MultiCoLA) of large community data sets. Nucleic Acids Res 38: e155.

Gramaje, D., Úrbez-Torres, J.R., and Sosnowski, M.R. (2018) Managing Grapevine Trunk Diseases With Respect to Etiology and Epidemiology: Current Strategies and Future Prospects. Plant Dis 102: 12–39.

Griffiths, S.M., Galambao, M., Rowntree, J., Goodhead, I., Hall, J., O’Brien, D., et al. (2020) Complex associations between cross-kingdom microbial endophytes and host genotype in ash dieback disease dynamics. J Ecol 108: 291–309.

Hardoim, P.R., van Overbeek, L.S., Berg, G., Pirttilä, A.M., Compant, S., Campisano, A., et al. (2015) The Hidden World within Plants: Ecological and Evolutionary Considerations for Defining Functioning of Microbial Endophytes. Microbiol Mol Biol Rev 79: 293–320.

Hofstetter, V., Buyck, B., Croll, D., Viret, O., Couloux, A., and Gindro, K. (2012) What if esca disease of grapevine were not a fungal disease? Fungal Divers 54: 51–67.

Hyde, K.D., Xu, J., Rapior, S., Jeewon, R., Lumyong, S., Niego, A.G.T., et al. (2019) The amazing potential of fungi: 50 ways we can exploit fungi industrially. Fungal Divers 97: 1–136.

Jousset, A., Bienhold, C., Chatzinotas, A., Gallien, L., Gobet, A., Kurm, V., et al. (2017) Where less may be more: how the rare biosphere pulls ecosystems strings. ISME J 11: 853–862.

Kenfaoui, J., Radouane, N., Mennani, M., Tahiri, A., El Ghadraoui, L., Belabess, Z., et al. (2022) A Panoramic View on Grapevine Trunk Diseases Threats: Case of Eutypa Dieback, Botryosphaeria Dieback, and Esca Disease. J Fungi (Basel) 8:.

Koch, H. and Schmid-Hempel, P. (2011) Socially transmitted gut microbiota protect bumble bees against an intestinal parasite. Proc Natl Acad Sci U S A 108: 19288–19292.

Kõljalg, U., Abarenkov, K., Nilsson, R.H., Larsson, K.-H., and Taylor, A.F.S. (2019) The UNITE Database for Molecular Identification and for Communicating Fungal Species. BISS 3: e37402.

Koskella, B., Meaden, S., Crowther, W.J., Leimu, R., and Metcalf, C.J.E. (2017) A signature of tree health? Shifts in the microbiome and the ecological drivers of horse chestnut bleeding canker disease. New Phytol 215: 737–746.

Lamichhane, J.R. and Venturi, V. (2015) Synergisms between microbial pathogens in plant disease complexes: a growing trend. Front Plant Sci 6: 385.

Larignon, P. and Dubos, B. (1997) Fungi associated with esca disease in grapevine. Eur J Plant Pathol 103: 147–157.

Larignon, P., Fontaine, F., Farine, S., Clément, C., and Bertsch, C. (2009) Esca et Black Dead Arm : deux acteurs majeurs des maladies du bois chez la Vigne. C R Biol 332: 765–783.

Lin, H. and Peddada, S.D. (2020) Analysis of compositions of microbiomes with bias correction. Nat Commun 11: 3514.

Lòpez-Fernàndez, S., Sonego, P., Moretto, M., Pancher, M., Engelen, K., Pertot, I., and Campisano, A. (2015) Whole-genome comparative analysis of virulence genes unveils similarities and differences between endophytes and other symbiotic bacteria. Front Microbiol 6: 419.

Lundberg, D.S., Lebeis, S.L., Paredes, S.H., Yourstone, S., Gehring, J., Malfatti, S., et al. (2012) Defining the core Arabidopsis thaliana root microbiome. Nature 488: 86–90.

Marchi, G., Peduto, F., Mugnai, L., Di Marco, S., Calzarano, F., and Surico, G. (2006) Some observations on the relationship of manifest and hidden esca to rainfall. Phytopathol Mediterr 45: S117–S126.

Markakis, E.A., Kavroulakis, N., Ntougias, S., Koubouris, G.C., Sergentani, C.K., and Ligoxigakis, E.K. (2017) Characterization of Fungi Associated With Wood Decay of Tree Species and Grapevine in Greece. Plant Dis 101: 1929–1940.

Martins, F., Mina, D., Pereira, J.A., and Baptista, P. (2021) Endophytic fungal community structure in olive orchards with high and low incidence of olive anthracnose. Sci Rep 11: 689.

Mendes, R., Kruijt, M., de Bruijn, I., Dekkers, E., van der Voort, M., Schneider, J.H.M., et al. (2011) Deciphering the rhizosphere microbiome for disease-suppressive bacteria. Science 332: 1097– 1100.

Millar, C.I. and Stephenson, N.L. (2015) Temperate forest health in an era of emerging megadisturbance. Science 349: 823–826.

Mina, D., Pereira, J.A., Lino-Neto, T., and Baptista, P. (2020) Impact of plant genotype and plant habitat in shaping bacterial pathobiome: a comparative study in olive tree. Sci Rep 10: 3475.

Mishra, S., Bhattacharjee, A., and Sharma, S. (2021) An Ecological Insight into the Multifaceted World of Plant-Endophyte Association. CRC Crit Rev Plant Sci 40: 127–146.

Monod, V., Hofstetter, V., Zufferey, V., Viret, O., Gindro, K., and Croll, D. (2022) Quantifying Trade-Offs in the Choice of Ribosomal Barcoding Markers for Fungal Amplicon Sequencing: a Case Study on the Grapevine Trunk Mycobiome. Microbiol Spectr 10: e0251322.

Monod, V., Zufferey, V., Wilhelm, M., Viret, O., Gindro, K., Croll, D., and Hofstetter, V. (2023) A systemic approach allows to identify the pedoclimatic conditions most critical in the susceptibility of a grapevine cultivar to esca/Botryosphaeria dieback. bioRxiv 2023.05.23.541976.

Mugnai, L., Graniti, A., and Surico, G. (1999) Esca (Black Measles) and Brown Wood-Streaking: Two Old and Elusive Diseases of Grapevines. Plant Dis 83: 404–418.

Pacifico, D., Squartini, A., Crucitti, D., Barizza, E., Lo Schiavo, F., Muresu, R., et al. (2019) The Role of the Endophytic Microbiome in the Grapevine Response to Environmental Triggers. Front Plant Sci 10: 1256.

Pedrós-Alió, C. (2006) Marine microbial diversity: can it be determined? Trends Microbiol 14: 257– 263.

Põlme, S., Abarenkov, K., Henrik Nilsson, R., Lindahl, B.D., Clemmensen, K.E., Kauserud, H., et al. (2020) FungalTraits: a user-friendly traits database of fungi and fungus-like stramenopiles. Fungal Divers 105: 1–16.

Pozo, M.J., Zabalgogeazcoa, I., Vazquez de Aldana, B.R., and Martinez-Medina, A. (2021) Untapping the potential of plant mycobiomes for applications in agriculture. Curr Opin Plant Biol 60: 102034.

Proença, D.N., Francisco, R., Kublik, S., Schöler, A., Vestergaard, G., Schloter, M., and Morais, P.V. (2017) The Microbiome of Endophytic, Wood Colonizing Bacteria from Pine Trees as Affected by Pine Wilt Disease. Sci Rep 7: 4205.

Rana, K.L., Kour, D., Sheikh, I., Yadav, N., Yadav, A.N., Kumar, V., et al. (2019) Biodiversity of Endophytic Fungi from Diverse Niches and Their Biotechnological Applications. In Advances in Endophytic Fungal Research: Present Status and Future Challenges. Singh, B.P. (ed). Cham: Springer International Publishing, pp. 105–144.

Reis, P., Magnin-Robert, M., Nascimento, T., Spagnolo, A., Abou-Mansour, E., Fioretti, C., et al. (2016) Reproducing Botryosphaeria Dieback Foliar Symptoms in a Simple Model System. Plant Dis 100: 1071–1079.

Rodriguez, R.J., White, J.F., Jr, Arnold, A.E., and Redman, R.S. (2009) Fungal endophytes: diversity and functional roles. New Phytol 182: 314–330.

Rolli, E., Marasco, R., Vigani, G., Ettoumi, B., Mapelli, F., Deangelis, M.L., et al. (2015) Improved plant resistance to drought is promoted by the root-associated microbiome as a water stress-dependent trait. Environ Microbiol 17: 316–331.

Romani, L. (2011) Immunity to fungal infections. Nat Rev Immunol 11: 275–288.

Romeralo, C., Martin-Garcia, J., Martinez-Alvarez, P., Jordan Munoz-Adalia, E., Goncalves, D.R., Torres, E., et al. (2022) Pine species determine fungal microbiome composition in a common garden experiment. Fungal Ecol 56:.

Saikkonen, K., Wäli, P., Helander, M., and Faeth, S.H. (2004) Evolution of endophyte-plant symbioses. Trends Plant Sci 9: 275–280.

Santini, A., Ghelardini, L., De Pace, C., Desprez-Loustau, M.L., Capretti, P., Chandelier, A., et al. (2013) Biogeographical patterns and determinants of invasion by forest pathogens in Europe. New Phytol 197: 238–250.

Säterberg, T., Jonsson, T., Yearsley, J., Berg, S., and Ebenman, B. (2019) A potential role for rare species in ecosystem dynamics. Sci Rep 9: 11107.

Segata, N., Izard, J., Waldron, L., Gevers, D., Miropolsky, L., Garrett, W.S., and Huttenhower, C. (2011) Metagenomic biomarker discovery and explanation. Genome Biol 12: R60.

Sheibani-Tezerji, R., Naveed, M., Jehl, M.-A., Sessitsch, A., Rattei, T., and Mitter, B. (2015) The genomes of closely related Pantoea ananatis maize seed endophytes having different effects on the host plant differ in secretion system genes and mobile genetic elements. Front Microbiol 6: 440.

Sieber, T.N. (2007) Endophytic fungi in forest trees: are they mutualists? Fungal Biol Rev 21: 75–89.

Singh, B.K., Delgado-Baquerizo, M., Egidi, E., Guirado, E., Leach, J.E., Liu, H., and Trivedi, P. (2023) Climate change impacts on plant pathogens, food security and paths forward. Nat Rev Microbiol 21: 640–656.

Sosnowski, M.R., Shtienberg, D., Creaser, M.L., Wicks, T.J., Lardner, R., and Scott, E.S. (2007) The influence of climate on foliar symptoms of eutypa dieback in grapevines. Phytopathology 97: 1284–1289.

Stewart, J.E., Kim, M.-S., Lalande, B., and Klopfenstein, N.B. (2021) Chapter 15 - Pathobiome and microbial communities associated with forest tree root diseases. In Forest Microbiology. Asiegbu, F.O. and Kovalchuk, A. (eds). Academic Press, pp. 277–292.

Surico, G. (2009) Towards a redefinition of the diseases within the esca complex of grapevine. 48: 5– 10.

Surico, G., Mugnai, L., and Marchi, G. (2006) Older and more recent observations on esca: a critical overview. Phytopathol Mediterr.

Tiew, P.Y., Mac Aogain, M., Ali, N.A.B.M., Thng, K.X., Goh, K., Lau, K.J.X., and Chotirmall, S.H. (2020) The Mycobiome in Health and Disease: Emerging Concepts, Methodologies and Challenges. Mycopathologia 185: 207–231.

Travadon, R., Rolshausen, P.E., Gubler, W.D., Cadle-Davidson, L., and Baumgartner, K. (2013) Susceptibility of Cultivated and Wild Vitis spp. to Wood Infection by Fungal Trunk Pathogens. Plant Dis 97: 1529–1536.

Trivedi, P., Leach, J.E., Tringe, S.G., Sa, T., and Singh, B.K. (2020) Plant-microbiome interactions: from community assembly to plant health. Nat Rev Microbiol 18: 607–621.

Turner, T.R., James, E.K., and Poole, P.S. (2013) The plant microbiome. Genome Biol 14: 209.

Vandenkoornhuyse, P., Quaiser, A., Duhamel, M., Le Van, A., and Dufresne, A. (2015) The importance of the microbiome of the plant holobiont. New Phytol 206: 1196–1206.

Vaz, A.B.M., Fonseca, P.L.C., Badotti, F., Skaltsas, D., Tomé, L.M.R., Silva, A.C., et al. (2018) A multiscale study of fungal endophyte communities of the foliar endosphere of native rubber trees in Eastern Amazon. Sci Rep 8: 16151.

Vieites, J.M., Guazzaroni, M.-E., Beloqui, A., Golyshin, P.N., and Ferrer, M. (2009) Metagenomics approaches in systems microbiology. FEMS Microbiol Rev 33: 236–255.

Wickham, H. ggplot2, Springer International Publishing.

Yadeta, K.A. and J Thomma, B.P.H. (2013) The xylem as battleground for plant hosts and vascular wilt pathogens. Front Plant Sci 4: 97.

